# Paracrine Signals from HIV-1 Infected Immune Cells Reprogram Cervical Cancer Pathways

**DOI:** 10.1101/2025.06.06.658239

**Authors:** Charles Ochieng’ Olwal, Ujjwal Rathore, Sara Makanani, Prashant Kaushal, Immy A. Ashley, Manisha R. Ummadi, Vincent Appiah, Alexandra Lindsey Djomkam Zune, Sophie Blanc, Declan Winters, Yennifer Delgado, Kapten Muthoka, Jacqueline M. Fabius, Manon Eckhardt, Robyn M. Kaake, Maureen Su, Oliver I. Fregoso, Judd F. Hultquist, Elkanah Omenge Orang’o, Danielle L. Swaney, George Boateng Kyei, Nevan J. Krogan, Peter Kojo Quashie, Yaw Bediako, Mehdi Bouhaddou

## Abstract

Persistent infection with human papillomavirus (HPV) is the primary cause of cervical cancer worldwide. Notably, women co-infected with HPV and human immunodeficiency virus type 1 (HIV-1) have a six-fold higher lifetime risk of developing cervical cancer compared to those without HIV, even when adhering to antiretroviral therapy (ART) and achieving T-cell reconstitution. While chronic HIV-1 infection is known to cause inflammation, how paracrine signals from immune cells alter signaling in cervical cells remain poorly understood. To address this, we conducted global transcriptomics analysis on cervical swabs from Kenyan women with HPV, stratified by HIV-1 and cancer status. Strikingly, women with HIV-1 showed cancer-like gene expression patterns in non-cancerous cervical epithelial cells. Complementary global mass spectrometry (MS) proteomics of cervical cells exposed to the secretome of HIV-1 infected primary CD4+ T-cells revealed altered expression of proteins in MAPK, PI3K-AKT, and β-catenin signaling pathways. Integrative network analyses of transcriptomic and proteomic datasets revealed that HIV-1 altered gene expression in key pathways known to drive cervical cancer, including genes commonly mutated in HIV-1-naïve disease. Notably, IRS-1, a key PI3K-AKT pathway activator, was found to be consistently upregulated in both participant samples and cell culture models, as were interferon-stimulated genes. Phosphoproteomics MS analysis confirmed PI3K-AKT pathway activation in cervical cells exposed to conditioned media from HIV-1-infected T-cells. Together, our findings uncover how HIV-1 reshapes cervical cell signaling via paracrine mechanisms and highlights the PI3K pathway as a potential therapeutic target in HIV-associated cervical cancer.

## Introduction

Cervical cancer, principally caused by high-risk human papillomavirus (HR-HPV) types, is the leading gynecological malignancy in human immunodeficiency virus type 1 (HIV-1) endemic regions, particularly in sub-Saharan Africa (SSA) (1,2). To accelerate cervical cancer elimination, the World Health Organization (WHO) proposed a ten-year triple intervention target in 2020. The intervention encompasses: (i) 90% HPV vaccination; (ii) 70% cervical screening coverage; and (iii) 90% access to treatment and care for precancerous cervical lesions (PCL) and invasive squamous cervical cell carcinoma (ICC) (3). Although massive roll-out of HPV vaccination programs in SSA could substantially reduce the incidences of cervical cancer, many young women and adolescent girls in SSA have already acquired some HR-HPV types (4,5) and may not benefit from the current HPV vaccines. Currently available HPV vaccines only protect against 9 out of 15 HR-HPV types (6) and their efficacy against established cancer is unknown. Infection with HIV-1 is associated with accelerated progression to cervical cancer, even in women with a fully reconstituted immune profile following adherence to antiretroviral therapy (ART) (7). Therefore, young women and adolescent girls in SSA, most of whom are highly vulnerable to HIV-1 infection (8,9), are at elevated risk of developing cervical cancer. In view of the shortcomings of the current HPV vaccines, and the fact that women in SSA mostly present at the late stages of cancer (10,11), effective treatment of ICC in the context of HIV/HPV coinfection is a critical component in the fight against this malignancy, warranting deeper investigation.

Women living with HIV-1 (WLWH) have a three-fold and six-fold elevated lifetime risk of developing PCL (12) and progressing to ICC (7), respectively, compared to those who have never acquired HIV-1. Increased prevalence of PCL (13,14) and its rapid progression to ICC have been attributed to HIV-associated CD4+ T-cell depletion (15,16). The occurrence and severity of other HIV/AIDS-defining cancers (*i*.*e*., Kaposi Sarcoma and non-Hodgins lymphoma) have declined dramatically following CD4 T-cell rebound upon adherence to ART. However, large population-based studies have reported stable incidences of cervical cancer even with highly effective ART (17–19). This suggests that accelerated progression to advanced cervical cancer in WLWH is not largely ascribed to immune suppression in these women. Thus, the molecular mechanism(s) that underlie accelerated progression to HPV-associated cervical cancer in WLWH is not fully understood. Deeper knowledge of these mechanisms could inform effective anti-cervical cancer interventions for WLWH.

Dysregulation of host gene expression is a hallmark of cancer development (20,21) and is linked with the activities of viral pathogens in several virus-associated cancers (22,23). While HPV directly infects cervical epithelial cells, HIV-1 does not, as cervical cells lack the receptor (*i*.*e*., CD4) and co-receptors (*i*.*e*., CCR5 and CXCR4) required for HIV-1 entry. However, the cervical epithelia are in close proximity to HIV-1 target immune cells, such as CD4+ T cells and macrophages, which are abundant in the cervico-vaginal environment (24). HIV-1 infection is associated with heightened systemic inflammation (25–28), which is linked with many HIV-related co-morbidities, including metabolic syndrome, neuropathic pain, chronic obstructive pulmonary disease, diabetes mellitus among others (29), irrespective of ART adherence. Furthermore, it has been shown that people living with HIV (PLWH) irreversibly and overly express certain host factors regardless of the duration of ART and levels of viral suppression relative to people who are HIV-naïve (30). We hypothesize that HIV-1 infection alters the secretome of immune cells, which alters pro-cancer pathways within proximal cervical cells. A thorough understanding of how HIV-1 infection remodels the transcriptomic and proteomic landscapes of human cervical cells via paracrine mechanisms could pinpoint the molecular mechanisms that HIV-1 infection alters to accelerate progression to advanced HPV-associated cervical cancer. In a previous network modeling study (31), we highlighted the phosphatidylinositol 3-kinase-AKT (PI3K-AKT) signaling pathway as converging between HIV-1-and HPV-host protein-protein interactions, potentially underlying enhanced cervical cancer development in HPV/HIV coinfection.

To test our hypothesis, we performed global transcriptomic profiling of cervical samples from HR-HPV infected black Kenyan women presenting with either a normal cervix (NC; cervix without lesions), PCL or ICC who were either HIV-infected or uninfected. In addition, we investigated the global changes in the abundance and phosphorylation of proteins in cervical cell cultures treated with the secretome of immune cells that were either infected with a replication-competent HIV-1 strain or uninfected as a control. Bioinformatics analysis and integration of data from the multiple system-level approaches suggested that HIV-1 infection activates and dysregulates pro-cancer signaling pathways, such as mitogen-activated protein kinase (MAPK) and PI3K-AKT, to accelerate HPV-associated cervical cancer development. Our findings uncover the molecular pathways potentially underlying accelerated progression to advanced HPV-associated cervical cancer in WLWH.

## Results

### Overview of participant characteristics and study design

In this study, we set out to understand the mechanisms underlying elevated progression to HPV-associated cervical cancer seen in WLWH. First, we carried out a cross-sectional study involving black Kenyan women from western Kenya, a region with high burden of cervical cancer (32) and HIV (33,34). We recruited women living with (n = 50) and without (n = 43) HIV-1 (Table S1) who were scheduled for cervical cancer screening by visual inspection with acetic acid (VIA), a simple and cost-effective cancer screening test commonly utilised in resource-limited settings such as Kenya (35). Upon providing written informed consent, the participants completed a questionnaire for medical, demographic and health histories to assess their eligibility for study enrollment. Prior to VIA, we collected two cervical swab samples from each participant who met the inclusion criteria (Table S2). Based on VIA screening results, we further stratified participants according to their cancer status: NC (*i*.*e*., those who were VIA negative), PCL (*i*.*e*., those with a VIA positive result), or ICC (*i*.*e*., those with a VIA positive or suspicious result and confirmed to have ICC by cervical biopsy test). All the HIV-positive participants were on ART and were not immunocompromised. For participants found to be infected with at least one of the International Agency for Research on Cancer (IARC) HR-HPV types (*i*.*e*., HPV16, HPV31, HPV33, HPV35, HPV52, HPV58, HPV18, HPV39, HPV45, HPV59, HPV51, HPV56, HPV53, HPV66, or HPV68) (36), the two cervical swab samples collected per participant were pooled followed by host RNA extraction and bulk RNA sequencing (RNA-seq; Figure 1A).

**Figure 1.**
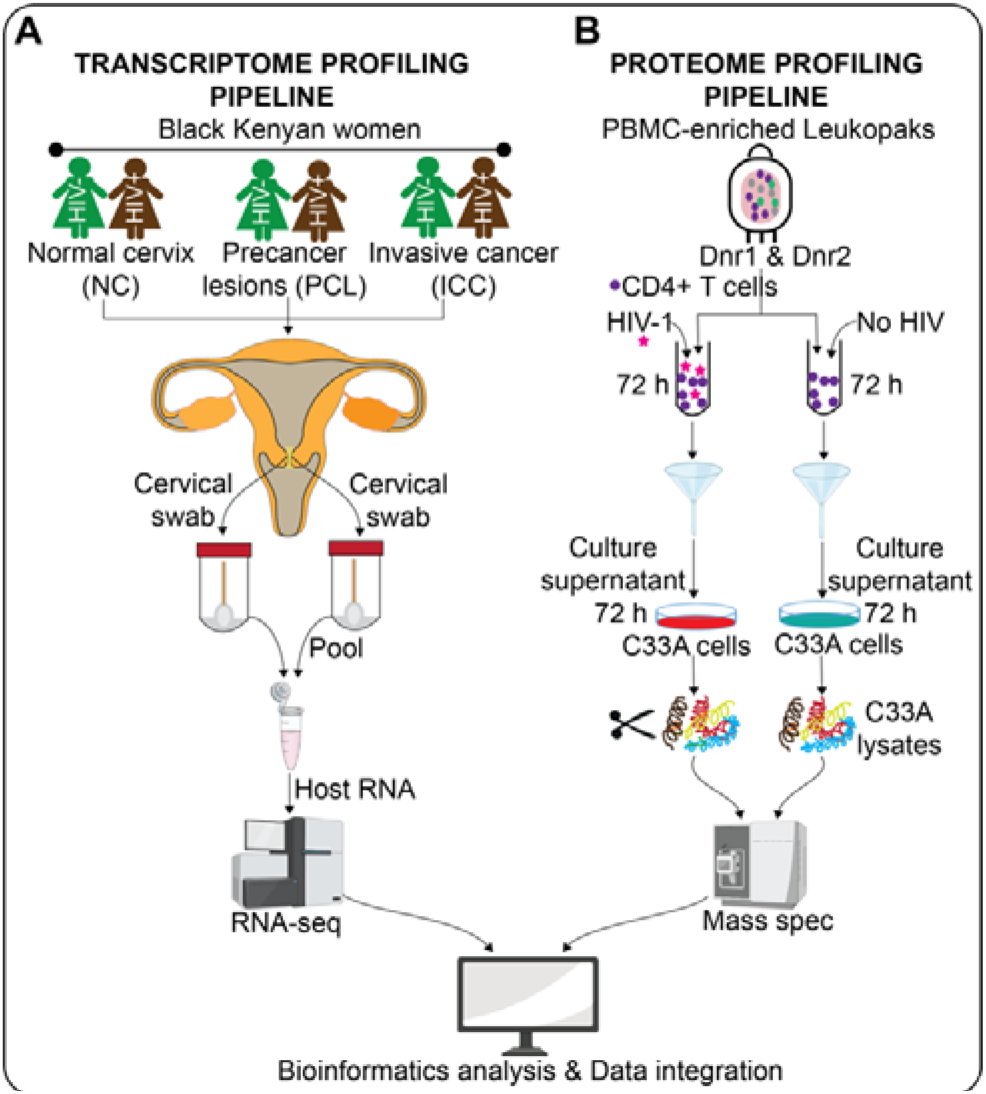
Summary of the study workflow. **(A)** Pairs of cervical swabs were collected from each of the HIV-infected or HIV-uninfected women. The two cervical swab samples per participant were pooled followed by PCR-based HR-HPV testing, host RNA extraction and RNA sequencing. **(B)** Primary CD4+ T-cells isolated from PBMC-enriched leukopaks from two healthy donors were infected with a replication-competent NL4-3 HIV-1 strain versus uninfected controls for 72 hours. Secretome of the CD4+ T-cells was used to stimulate C33A cells for 72 hours. After 72 hours, the C33A cell lysates were digested and subjected to global quantitative mass spectrometry (MS) based proteomic analysis. Transcriptomics and proteomics outputs were analysed independently and integrated using bioinformatics approaches.

Since human cervix is comprised of immune cells known to harbor HIV (24), we next set out to understand how the secretome of HIV-infected immune cells impact the proteomic landscape of cervical cells. To this end, we cultured an HPV-negative human cervical cancer cell line [*i*.*e*., C33A; (37)] with the media from primary CD4+ T-cells infected with a replication-competent HIV-1 NL4-3 strain versus uninfected controls (Figure S1). Briefly, bulk CD4+ T-cells were isolated from peripheral blood mononuclear cells (PBMCs)-enriched leukapheresis products (leukopaks) from two healthy donors by magnetic negative selection (*i*.*e*., to avoid activation) and challenged with HIV-1 NL4-3 for 72 hours prior to the collection of conditioned media. C33A cells were cultured with the conditioned media for 72 hours. We then quantified global changes in protein abundance in the cervical cells using bottom-up mass spectrometry (MS) proteomics (Figure 1B). Finally, we performed independent and integrated bioinformatics analyses of the transcriptomic and proteomic datasets (Figure 1A-B) to identify the cervical epithelial signaling pathways perturbed in the context of HIV-1 infection.

### HIV-1 infection evokes a cancer-like state in the normal cervix

We first sought to understand how HIV-1 infection altered the transcriptomic landscape between NC and ICC participants in the context of HR-HPV coinfection. As mentioned above, our clinical study recruited 50 women with HIV-1 and 43 without HIV, who were determined to either have NC, PCL or ICC (Figure 2A). Upon testing for HR-HPV positivity, we found 57% of NC participants were HPV positive and all participants with PCL or ICC were HPV positive (Figure 2B), showing that cervical cancer in our study population is largely associated with HPV. We performed global transcriptomics (*i*.*e*., RNA-seq) analysis on cervical swabs from women who were all HR-HPV positive but split based on HIV-1 positivity (Figure 2C). The participants (n = 35) selected for transcriptome profiling were between the age of 22 and 50 years (Figure 2D). We noted that the median ages of HIV-infected participants with PCL (*i*.*e*., 37) or ICC (*i*.*e*., 35) were relatively lower compared to that of HIV-uninfected women with PCL (*i*.*e*., 45) or ICC (*i*.*e*., 43) (Figure 2D), which suggest that WLWH are likely to develop precancer or invasive cervical cancer at a younger age compared to women without HIV.

**Figure 2.**
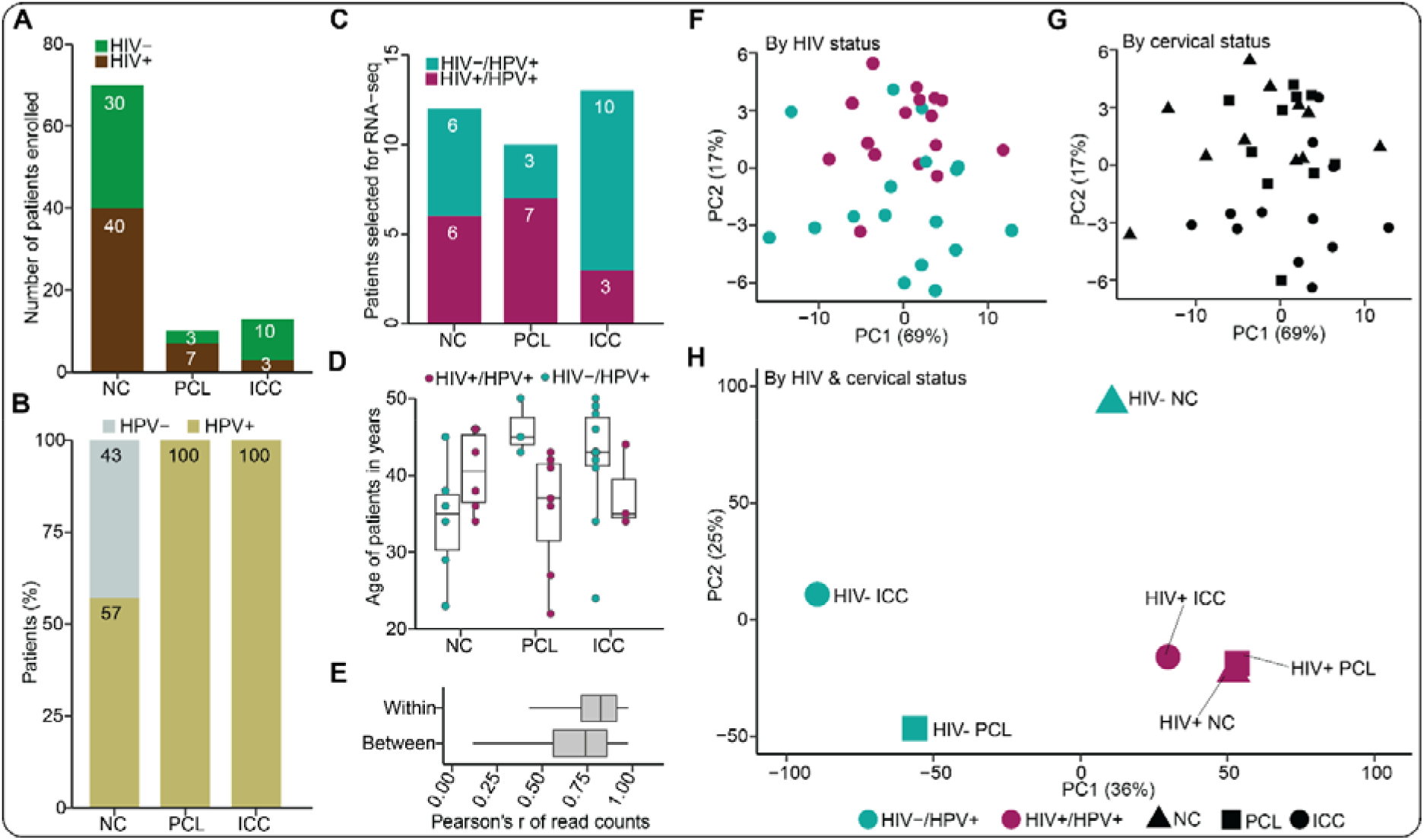
Transcriptome profiles of cervical samples depict differences between HIV-infected and uninfected women. **(A)** The distribution of 93 participants recruited into the study based on their HIV and cervical status. **(B)** The distribution of HR-HPV infection across the three cervical statuses as determined by a PCR-based genotyping technique. **(C)** The distribution of HR-HPV positive participants and HIV-infected or uninfected participants across the three cervical statuses who were selected for RNA-seq analysis. **(D)** The age distribution of HR-HPV positive and HIV-infected or HIV-negative participants across the three cervical statuses selected for RNA-seq. Each dot represent a participant. The center line in each box depicts the median whereas the lower and upper edge of each box represent the 25th and 75th percentile values, respectively. The whiskers represent 1.5 times the interquartile range. **(E)** The correlation of feature counts within and between participant samples. The center line in each box represents the median whereas the lower and upper edge of each box depict the 25th and 75th percentile values, respectively. The whiskers represent 1.5 times the interquartile range. (**F-G**) Computational principal component analysis (PCA) plots of the feature counts from individual participants based on either their HIV status or cervical status. (**H**) A PCA plot of feature counts averaged per HIV and cervical status.

Next, we assessed whether the transcriptomic profiles could distinguish participants based on their HIV and/or cervical cancer status. First, we decided to assess the intra-and inter-sample correlation. We observed high correlations within and between participant samples (correlation coefficients > 0.7; Figure 2E). Computational principal component analysis (PCA) of the feature counts data revealed a separation of individual participants with regards to HIV (Figure 2F) or cervical cancer status (Figure 2G) along PC2. We then performed PCA on feature counts averaged across participants grouped according to HIV and cervical cancer status. For HIV-negative women, the PCA revealed a clear segregation across cervical cancer stage (Figure 2H). Interestingly, HIV-positive women who had NC, PCL or ICC clustered together (Figure 2H). This latter observation suggests that HIV-1 infection evokes a cancer-like state in the NC that persists and predisposes these women to PCL and ICC via dysregulated signaling mechanisms. Our observations further suggest that HIV-1 infection contributes to the entire continuum of HPV-associated cervical cancer progression, that is, from initiation to advanced stages.

### Genes upregulated in the cervix of WLWH are enriched in pro-cancer pathways

We next sought to map differentially expressed genes (DEGs) to molecular pathways, comparing them across HIV and cancer status. Our analysis revealed 884, 50, and 122 genes upregulated in the HIV-positive group, relative to the HIV-negative group, for NC, PCL, and ICC, respectively (Figure 3A). Very few genes were found to be downregulated. Intriguingly, participants with NC displayed the largest differences based on HIV status whereas a much smaller magnitude of change was observed for participants with PCL or ICC (Figure 3A). This observation further supports the idea that HIV induces a cancer-like state in normal cervical epithelial cells, which becomes less distinguishable once cancer has developed (*i*.*e*., PCL or ICC). Since NC clustered with PCL and ICC for WLWH (Figure 2H) and HIV status was associated with marked transcriptional changes within the NC group (Figure 3A), we decided to focus on delineating the pathways that were dysregulated by HIV in the NC group.

**Figure 3.**
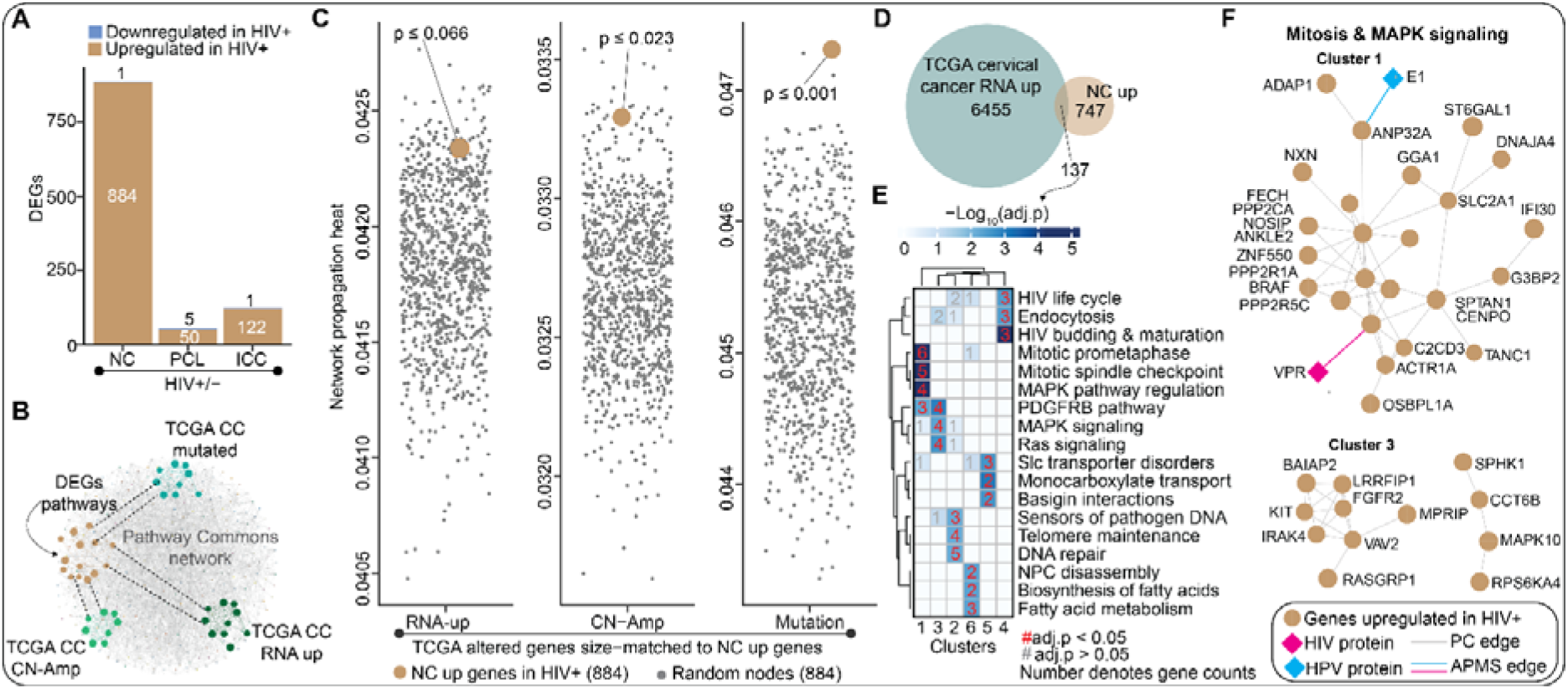
Transcriptome profiling of participant samples identifies cancer-related genes perturbed by HIV-1 infection. **(A)** Bar plots summarizing the number of differentially expressed genes (DEGs) in HIV-infected women across different cervical statuses. The DEGs were based on a cutoff of |log_2_FC| > 2 and p < 0.05. **(B)** Cartoon illustrating the closeness of the molecular pathways associated with the DEGs and known pro-cervical cancer genes (*i*.*e*., genes commonly altered in TCGA cervical cancer Pancancer cohort) within a larger biological network such as Pathway Commons network. **(C)** Jitter plots comparing the median network propagation heats for the 884 genes upregulated in HIV-infected women with NC versus the medians for size-matched random nodes sampled 1000 times from each of the network propagation outputs. Separate diffuse heat network propagation of the top 884 mRNA up, CN-amplified or mutated genes from TCGA cervical cancer Pancancer cohort was performed using Pathway Commons for base network. The median propagation heat of the 884 NC upregulated genes and that of 884 random nodes sampled 1000 times were plotted per network propagation. The p-value of the NC up genes per genetic alteration was calculated by first ranking the median heats then dividing the position of the NC up genes by the number of randomizations (*i*.*e*., 1000 times). Network propagation outputs are provided in Table S3. **(D)** Venn diagram showing the number of genes upregulated in both the HIV-positive NC group and participants listed in the TCGA cervical cancer Pancancer study cohort. **(E)** Heatmap showing the top three canonical pathways associated with each of the subnetwork clusters from a network generated from overlapping genes in (D) above. The gene-set overrepresentation analysis was performed for the network subclusters. The p-values were calculated by hypergeometric test with multiple hypothesis testing correction (false discovery rate; FDR). A complete set of enriched biological pathways is provided in Table S4. **(F)** The subnetwork clusters associated with pro-cancer pathways, such as mitosis and MAPK signaling, are highlighted. A complete network with all the six subnetwork clusters is shown in Figure S2.

To assess whether the genes upregulated in the WLWH-NC group were relevant to cervical cancer development, we used a network modeling approach to evaluate whether these genes shared molecular networks with genes from The Cancer Genome Atlas (TCGA) cervical cell carcinoma database that were either (i) transcriptionally upregulated (“RNA-up”; n = 884), (ii) copy number amplified (“CN-Amp”; n = 884), (iii) or highly (n = 884) mutated (“Mutation”; Figure 3B). Specifically, we used network propagation, a method that simulates “heat diffusion” across a molecular interaction network to assess the functional proximity between gene sets. We applied this approach using the Pathway Commons network, initiating propagation from the three sets of genes extracted from TCGA cervical cancer database. To quantify the connectivity between our gene set and known cancer-related genes, we measured the median “heat” accumulated on genes associated with RNA upregulation (RNA-up), copy number amplification (CN-Amp), or mutation in cervical cancer TCGA datasets. We then compared these values to distributions generated from 1,000 random gene sets matched for size, allowing us to calculate empirical p-values for each gene category. Strikingly, in each scenario, our set of 884 genes were highly ranked relative to the random sets of genes (Figure 3C). Our results suggest genes upregulated by HIV in NC individuals share molecular networks with genes that are commonly altered in cervical cancer participants.

Next, we sought to hone into the molecular pathways shared between the participants in our study and those from TCGA. We identified 137 genes that directly overlapped between RNA-up genes (n = 6455) from TCGA cervical cancer participants and from this study (n = 884; Figure 3D). We extracted an interconnected subnetwork of these 137 genes from the Pathway Commons network and performed hierarchical clustering to identify 6 highly interconnected sub-clusters (Figure S2 and 3E). A gene set overrepresentation analysis (GSOA) (31) using canonical pathway terms revealed HIV-related gene pathways, endocytosis, mitosis/cell cycle, Ras/ERK, DNA damage/repair, and pathogen sensors indicative of innate immune regulation (Figure 3E). Intriguingly, closer inspection of clusters 1 and 3 revealed the genes in these pathways and two of them, *ANP23A* and *DYNC1H1*, were previously known (38,39) to physically interact with HPV E1 and HIV-1 Vpr, respectively (Figure 3F). Overall, our results suggest that HIV-1 infection regulates pro-cancer pathways in normal cervical cells.

### Secretome of HIV-1 infected immune cells regulates pro-cancer proteins in cervical cells

Next, we sought to evaluate how paracrine signals from HIV-infected CD4+ T-cells remodel the proteome of cervical epithelial cells in culture. To assess this, we set up a controlled experimental system in which we quantified protein abundance changes in cervical cells treated with conditioned media from primary CD4+ T-cells infected or uninfected with a replication-competent HIV-1 strain. Specifically, we isolated CD4+ T-cells from participant-derived PBMCs from two donors, isolated CD4+ T-cells by magnetic negative selection, infected them with NL-43 HIV-1 for 72 hours, and harvested the conditioned cell culture media. We then cultured C33A cells in the CD4 T-cell conditioned media for 72 hours before lysing the cells and performing global mass spectrometry abundance proteomics.

Our analysis revealed altered expression of several endogenous proteins in cervical cells upon treatment with T-cell conditioned media, with a modest degree of variability by T-cell donor (Figure 4A). For uniformity with the transcriptomic analysis, we focused on upregulated proteins in the cervical cell for this analysis. Like in the preceding section, we decided to assess whether the 460 upregulated cervical cell proteins (union of T-cell donors 1 and 2) were relevant to cervical cancer development. To this end, we assessed if the proteins upregulated in the cervical cells are in the same network space as genes already known to be associated with cervical cancer development (Figure 4B), which we defined as proteins known to be proteomically upregulated, undergo CN-Amp, or are highly mutated in the squamous cervical cell carcinoma participants from TCGA. Using Pathway Commons for the base network, we performed three separate diffused heat network propagations with the top upregulated proteins (n = 100; all the proteins upregulated in the cervical cancer TCGA cohort), CN-Amp (n = 460), or mutated genes (n = 460; Figure 4B). As in the transcriptomics analysis, our results showed that the median propagation heat for the 460 proteins upregulated in cervical cells was higher than the median heat for 460 other random genes sampled 1000 times from each of the three network propagation outputs (Figure 4C). Our results suggest that proteins upregulated by the secretome of HIV-1 infected immune cells are network proximal to known cervical cancer genes pulled from the cervical cancer TCGA dataset, suggesting that the HIV-induced secretome of the immune cells upregulates cervical proteins that are most likely implicated in cervical cancer development.

**Figure 4.**
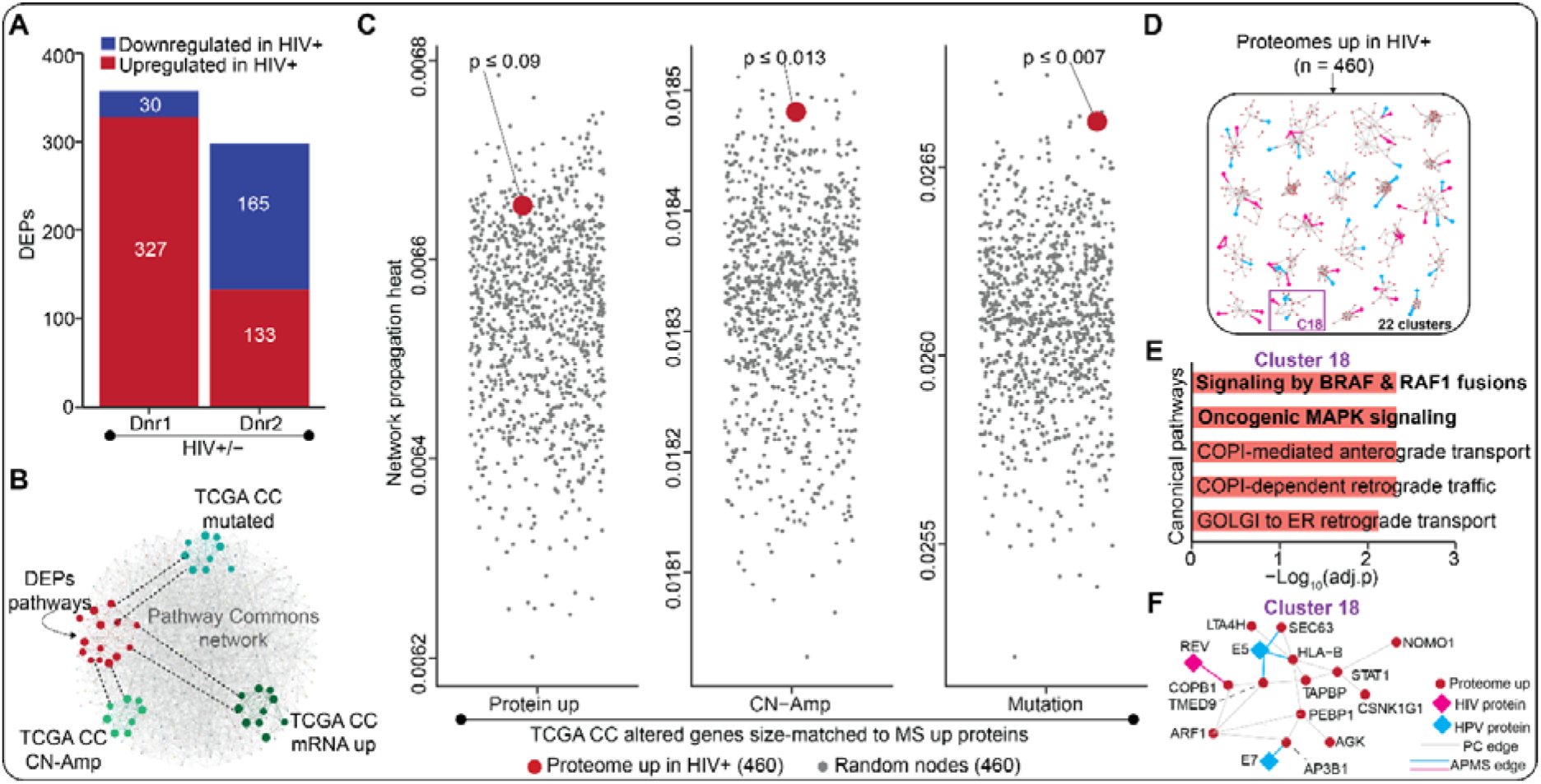
Proteomic profiling identifies pro-cancer pathways impacted by the secretome of HIV-1 infected immune cells. **(A)** Bar plots showing the number of differentially expressed proteins (DEP) in cervical cells stimulated with secretome of HIV-infected primary CD4+ T-cells isolated from two healthy donors compared to uninfected controls. The DEPs were determined based on |log2FC| > 1 and p < 0.05 cutoffs. **(B)** Cartoon illustrating the closeness of the molecular pathways associated with the DEPs and known pro-cervical cancer genes (*i*.*e*., genes commonly altered in TCGA cervical cancer Pancancer cohort) within a larger biological network such as Pathway Commons network. **(C)** Jitter plots comparing the median network propagation heats for the 460 proteins upregulated in cervical cells stimulated with secretome of HIV-infected immune cells versus the medians for size-matched random nodes sampled 1000 times from each of the network propagation outputs. Separate diffuse heat network propagation of the top 460 protein up, CN-Amp or mutated genes from TCGA cervical cancer Pancancer cohort was performed using Pathway Commons for base network. The median network propagation heat of the 460 upregulated proteins and that of 460 random nodes sampled 1000 times were plotted for each of the three network propagations performed. The p-value of the upregulated cervical cell proteins per genetic alteration was calculated by first ranking the median heats then dividing the position of the upregulated proteins by the number of randomizations (*i*.*e*., 1000 times). Network propagation outputs are provided in Table S5. **(D)** A dummy biological network generated from the upregulated proteins using Pathway Commons for base network. The network was clustered into 22 subnetworks. See Figure S3 for the actual network. **(E)** Bar plots highlighting the top five canonical pathways associated with cluster 18 of the network in (**D**). This bar plot was obtained from a GSOA performed on the subnetwork clusters from D. The p-values were calculated by hypergeometric test with multiple hypothesis testing correction (FDR). An enrichment heatmap highlighting the top three canonical pathway terms associated with all the 22 subnetwork clusters is provided as Figure S4. A complete set of enriched biological pathways are provided in Table S6. **(F)** The subnetwork cluster associated with MAPK signaling extracted from the bigger network in Figure S3 annotated from Pathway Commons network.

We next sought to identify the molecular pathways associated with the cervical cell proteins upregulated by the secretome of the HIV-infected immune cells. As above, we extracted a subnetwork from Pathway Commons for the proteins upregulated in cervical cells treated with immune cell secretome into a network and clustered it into 22 subnetwork clusters (Figure 4D and Figure S3). In line with our transcriptomics data, GSOA on these 22 subnetwork clusters showed that cluster 18 is enriched in MAPK signaling (Figure 4E-F). Altogether, these findings suggest that HIV-induced immune cell secretome alters pro-cancer pathways within cervical cells.

### Integration of mRNA and protein data highlights beta-catenin and MAPK as key driver pathways

We next sought to integrate the transcriptomics data with the proteomics data to ask if there are molecular pathways altered by HIV-1 infection at both the mRNA and protein levels. To identify pathways converging between the two datasets, we performed GSOA on the 460 proteins upregulated in cervical cells by the secretome of HIV-infected immune cells and size-matched top mRNAs upregulated in WLWH with NC. Our analysis revealed that the transcriptome and proteome data converge on three pro-cancer pathways: cell cycle (*i*.*e*., mitosis), the beta-catenin pathway, and “diseases of signal transduction” (Figure 5A). To gain a more granular picture of the “diseases of signal transduction” term, we assessed the log2FC of the 39 genes in this category found across the transcriptomic and proteomic datasets (Figure 5B). Although some genes were upregulated in both datasets (*e*.*g*., *IRS1, SMAD4, SKP1*, and *STAT1*) others were not (*e*.*g*., *PSMD5, PSMA6, PSMB4*, and *ATG7*). However, even though we observed examples of disjointed regulation at the gene level, we found overlapping regulation at the pathway level. To have a better resolution of the molecular pathways the 39 genes are implicated in, we visualized the genes in a STRING network and clustered it into three subnetworks followed by GSOA. In alignment with the independent enrichment analysis of transcriptome and proteome datasets, GSOA of the subnetwork clusters of genes associated with “diseases of signal transduction” term also showed enrichment in MAPK signaling pathway (Cluster 1; Figure 5C). To this subnetwork, we added HIV and HPV viral proteins based on previously discovered HIV-and HPV-host protein-protein interactions from affinity purification-MS experiments (38,39) (Figure 5C). We noted that the molecular networks associated with the diseases of signaling transduction were hijacked by HIV and HPV proteins (Figure 5D), implying that HIV-1 and HPV proteins are directly involved in the dysregulation of these pathways in cervical cells.

**Figure 5.**
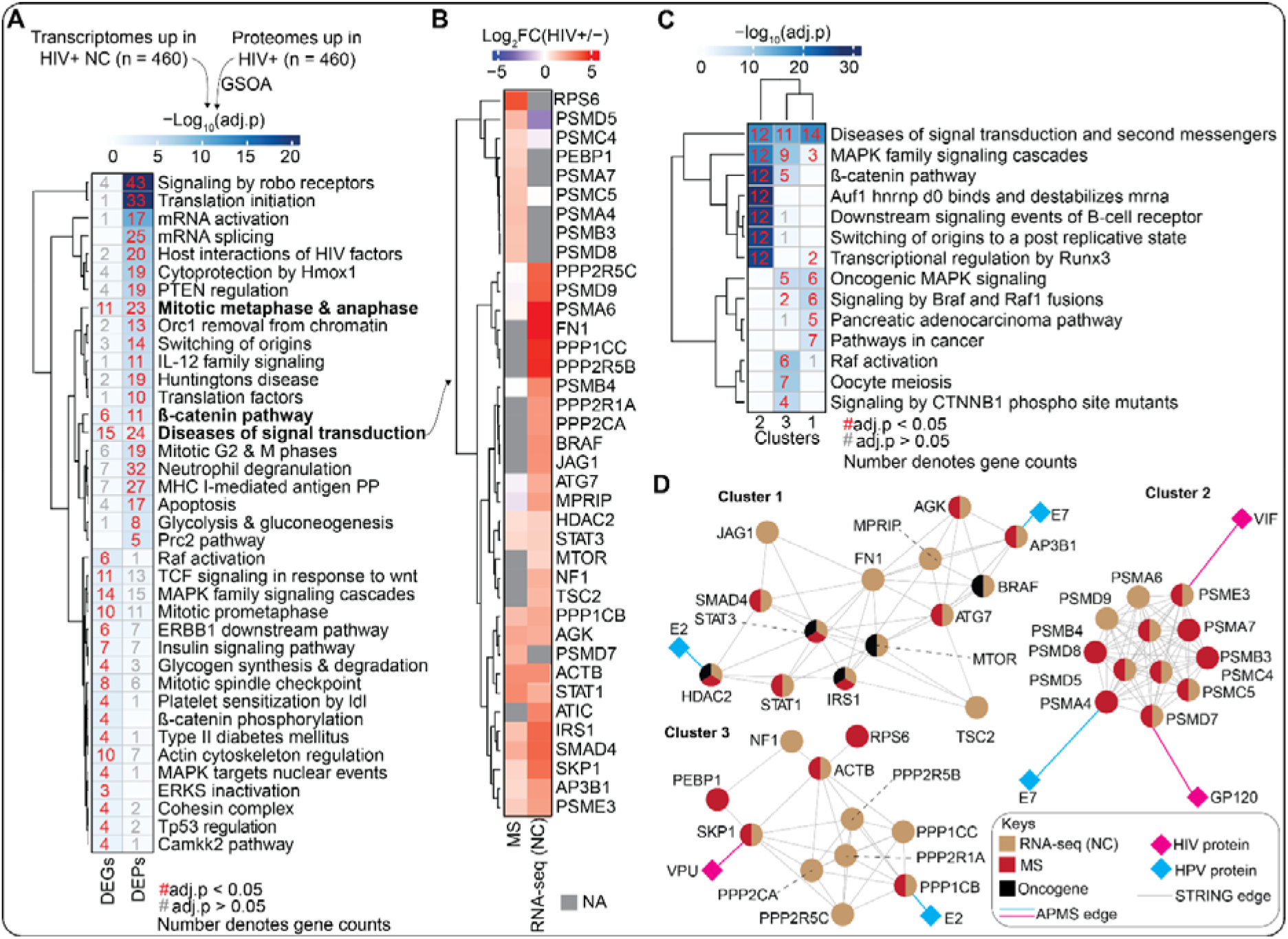
HIV-1 infection alters pro-cancer signaling pathways at the gene and protein levels. **(A)** Heatmap highlighting the canonical pathways from GSOA of significantly upregulated proteins in cervical cells treated with secretome of immune cell infected with HIV-1 and size-matched upregulated genes in WLWH with NC from proteomics and transcriptome profiling, respectively. The numbers depict count of proteins mapping to each canonical pathway term. Red numbers indicate a significant (adj.p value < 0.05) enrichment whereas gray numbers indicate a non-significant enrichment. Background color in the heatmap denotes the −log_10_adj.p values as shown on color bar. The p-values were calculated by hypergeometric test with multiple hypothesis testing correction (FDR). A complete set of enriched biological pathways are provided in Table S7. **(B)** A heatmap comparing the log2FC of the genes enriched for diseases of signal transduction in both the transcriptome and proteome data. **(C)** Heatmap depicting the canonical pathways from GSOA of three subnetworks annotated from STRING network using genes enriched for diseases of signal transduction in (B). The numbers depict count of proteins mapping to each canonical pathway term. Red numbers indicate a significant (adj.p value < 0.05) enrichment whereas gray numbers indicate a non-significant enrichment. Background color in the heatmap denotes the −log_10_adj.p values as shown on color bar). The p-values were calculated by hypergeometric test with multiple hypothesis testing correction (FDR). A complete set of enriched biological pathways are provided in Table S8. **(D)** Subnetworks of genes enriched for diseases of signal transduction in (B and C) annotated from STRING network. The HIV and HPV proteins were added based on viral-host protein-protein interactions generated in previous affinity purification MS studies (38,39).

### HIV-1 induced paracrine signals from immune cells activate the PI3K-AKT pathway in cervical cells

Our transcriptome and proteome studies identified molecular pathways that are likely dysregulated by HIV-1 infection to accelerate progression to HPV-associated cervical cancer. However, the two omics approaches cannot precisely tell whether a molecular pathway is activated or inactivated. Protein phosphorylation is an important regulator of cellular processes and plays a critical role in intracellular signal transduction (40). Thus, to clarify whether HIV-1 infection activates or inactivates pro-cancer pathways in cervical cells, we interrogated the global phosphorylation landscape of cells exposed to secretome of primary CD4+ T-cells infected with a replication-competent HIV-1 strain and uninfected controls similar to the workflow above for abundance proteomics (Figure 1B). Our analysis revealed that the secretome of HIV-infected immune cells differentially regulates phosphorylation of proteins in cervical cells (Figure 6A-B). Phosphorylation signaling is largely governed by protein kinases; hence, we used The Kinase Library, an algorithm of experimentally derived kinase sequence motifs, to infer kinase activities from the phosphoproteomics data (41,42). This analysis revealed activation of several kinases associated with MAPK and PI3K-AKT signaling, including AKT1, AKT2, AKT3, NLK, JNK1/MAPK8, and SMG1 (Figure 6C), implying that HIV-induced secretome activates the two pathways in cervical cells. Interestingly, we observed that insulin receptor substrate 1 (IRS1) is significantly expressed in both the transcriptomic and proteomic data (Figure 6D). Increased expression of IRS1 has been associated with the activation of both MAPK and PI3K signaling pathways (43,44). In addition, using decoupleR v2.12.0 package, we computationally investigated the transcription factor activity of genes upregulated in WLWH with NC inferred from prior knowledge (45). This analysis revealed that CTNNB1 is the topmost activated transcription factor in HIV-1 infected women with NC relative to those who are HIV naïve (Figure 6E).

**Figure 6.**
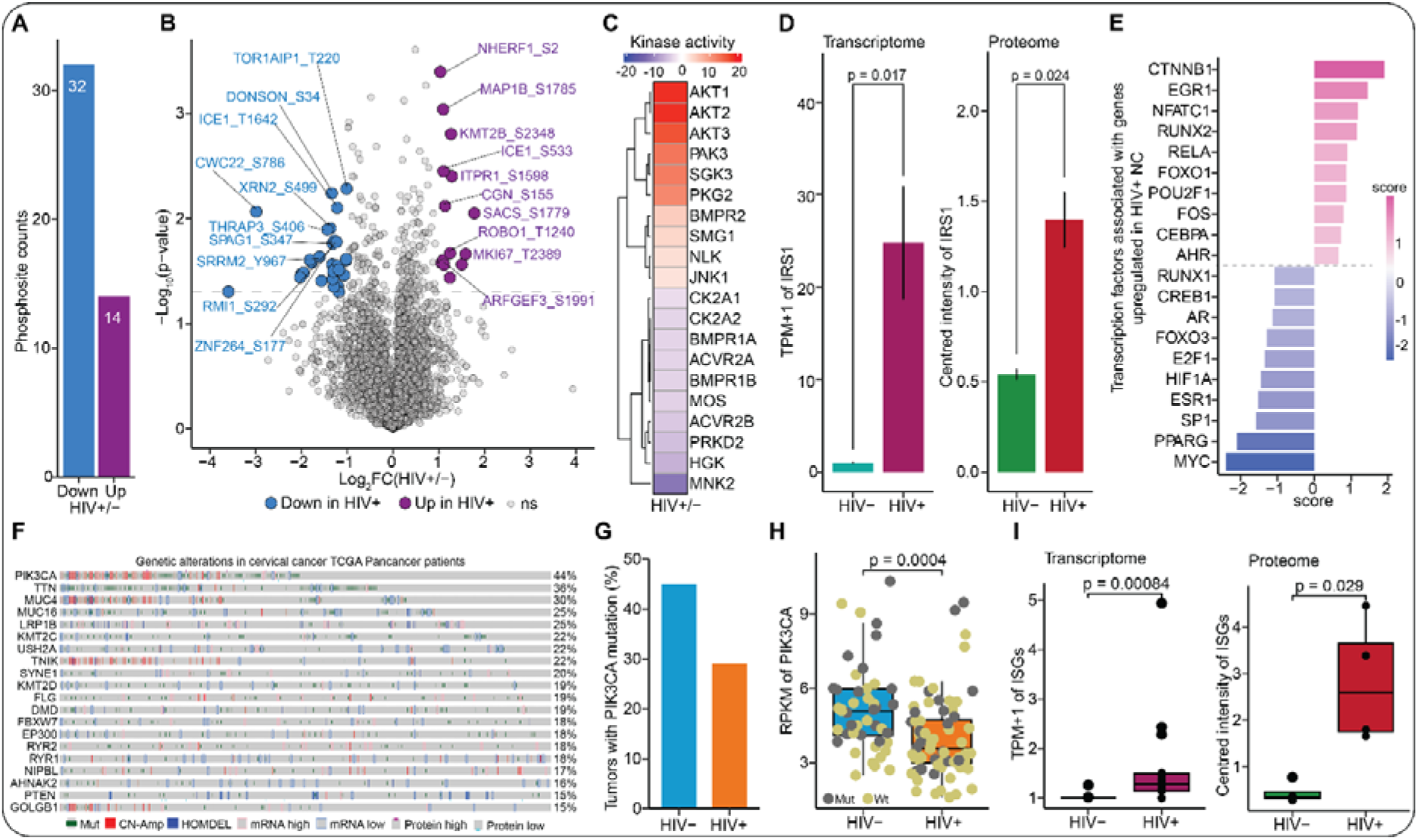
HIV-1 infection activates the PI3K signaling pathway. **(A)** Bar plots showing the number of differentially changing phosphosites in cervical cells treated with secretome of HIV-infected or uninfected primary CD4+ T-cells. Differential changes in the phosphosites were determined based on|log2FC| > 1 and p < 0.05 cutoffs. **(B)** Volcano plots of phosphoproteomics highlighting changes in phosphosites between cervical cells treated with secretome of HIV-1 infected and uninfected primary CD4+ T-cells. The colored dots depict the differentially changing phosphosites at |log2FC| > 1 and p < 0.05 cutoffs. **(C)** Heatmap showing the kinase activities of phosphorylated substrates from global MS phosphoproteomics analysis of cervical cells exposed to secretome of HIV-infected or uninfected primary CD4+ T-cells. The colors indicate an increase (red) or decrease (blue) in kinase activity. **(D)** Bar plots displaying the normalized gene expression (*i*.*e*., transcripts per million + 1) and protein expression levels (mean-centered intensities) of IRS1. Statistical comparison was performed using Wilcoxon sum ranked test. **(E)** Bar plots ranking the transcription factors activated or deactivated by the genes upregulated in WLWH with NC relative to their HIV-counterparts. **(F)** Oncoplot showing the alteration frequencies from the cervical cancer (TCGA Pancancer) dataset for the top 10 mutated genes in cervical cancer (cBioPortal). Each vertical gray column represents a participant and various genetic alterations of genes in each participant are indicated in the color legend. **(G)** Bar plots summarizing the proportion of HIV-positive and HIV-negative Ugandan cervical cancer tumors with PIK3CA mutation. Data from Gagliardi and colleagues (46). **(H)** Box plots depicting expression (RPKM) of mutant or wild-type PIK3CA gene in HIV-positive and negative cervical cancer tumors from Ugandan participants. The center line in each box depicts the median whereas the lower and upper edge of each box represent the 25th and 75th percentile values, respectively. The whiskers represent 1.5 times the interquartile range. Each dot represents a tumor. Yellow and dark gray dots represent tumors with wild-type and mutant PIK3CA. Statistical comparison was performed using Wilcoxon sum ranked test. Data from Gagliardi and colleagues (46) **(I)** Box plots showing the gene (TPM+1) and protein expression levels (*i*.*e*., mean-centered intensities) of known interferon-stimulated genes. The center line in each box depicts the median whereas the lower and upper edge of each box represent the 25th and 75th percentile values, respectively. The whiskers represent 1.5 times the interquartile range. Each dot represents an ISG. Statistical comparison was performed using Wilcoxon sum ranked test.

Our finding that HIV-1 infection activates PI3K signaling pathway is corroborated by an analysis of data extracted from TCGA cervical cancer participants, which ranked PIK3CA, the major catalytic subunit of PI3K pathway, as the topmost altered gene (Figure 6F). Interestingly, another study from a Ugandan cervical cancer cohort reported that mutation (Figure 6G) and expression (Figure 6H) of the PIK3CA gene was markedly lower in WLWH compared to women without HIV (46). This implies that HIV infection may phenocopy PI3K pathway mutations observed in HIV-negative contexts. Since our previous study (31) had shown that inflammatory factors can activate PI3K-AKT pathway, we asked whether exposed cervical cells experienced inflammation. We quantified the expression of interferon-stimulated genes (ISGs (47,48) in our transcriptomics and proteomics datasets and found significantly higher expression of a number of ISGs in HIV-infected compared to the HIV-uninfected conditions (Figure 6I and Figure S5), suggesting that oncogenic and interferon signaling are co-occurring. Altogether, our results suggest that the paracrine signals from HIV-1 infected immune cells activate the PI3K-AKT signaling pathway, upregulate IRS1, activate CTNNB1 transcription factor activity, and induce inflammatory signaling, all of which may accelerate the development of advanced cancer phenotypes.

## Discussion

How HIV-1 infection contributes to the acceleration of HPV-associated cervical cancer remains a mystery. The present study sought to address this question using multiple system-level approaches performed on cervical samples obtained from HPV-positive and HIV-infected or HIV-uninfected Black Kenyan women with NC, PCL or ICC. In parallel, cervical cells in culture were stimulated with the secretome of HIV-infected or uninfected immune cells.

First, we found that all participants with PCL and ICC were HR-HPV positive, confirming that cervical cancer in our study setting is largely ascribed to HR-HPV, as expected. Furthermore, HIV-positive women with PCL or ICC had considerably lower median age relative to those without HIV. This observation is in agreement with previous reports (49,50) and suggests that WLWH are predisposed to precancer and cervical cancer at a younger age compared to the HIV-naïve ones. Surprisingly, a PCA of read counts averaged per HIV and cervical status revealed that HIV-infected NC, PCL and ICC participants clustered together. This outcome suggests that HIV-1 infection evokes a cancer-like microenvironment in the HPV-positive normal cervix, which persists and predisposes women to accelerated progression to advanced cervical cancer via signaling mechanisms. Our data support the hypothesis that HIV-1 infection contributes to the early stages of HPV-associated cervical cancer development, supported by the observation that most transcriptomic changes occurred in the HIV-positive NC group, suggesting that HIV-1 may introduce irreversible changes in cervical cells at early stages. Additionally, the clustering of HIV-positive NC, PCL, and ICC samples implies a shared HIV-driven transcriptional program across disease stages. These findings may help explain the accelerated progression to cervical cancer observed in WLWH.

The present study sought to identify the molecular pathways that are targeted by HIV-1 infection during cervical cancer development. Functional enrichment of both transcriptome and proteome data highlighted that HIV-1 infection upregulates the expression of several cancer-associated pathways in cervical cells, including MAPK, cell cycle, DNA repair, RAS and WNT and PI3K signaling. The role of MAPK, cell cycle, RAS, DNA repair and PI3K pathways in cancer development is well documented (51,52). PIK3CA, the largest catalytic subunit of PI3K, is the second most altered gene in cervical cancer participants listed in cBioPortal and has emerged as the topmost mutated gene in cervical cancer participants across ethnicities (46,53–56). It is noteworthy that the dysregulation of these molecular pathways occurred in the normal cervix and were also dysregulated by secretome of HIV-1 infected immune cells. These findings suggest that HIV-1 infection of immune cells within the cervico-vaginal environment impacts major pro-cancer pathways in cervical cells and could explain why WLWH are prone to PCL and progression to advanced HPV-associated cervical cancer relative to HIV-naïve women. However, it remains to be determined whether the molecular pathways highlighted above are cooperatively targeted by both HIV-1 and HPV infection or not. Future studies would be needed to identify the nature of cooperation by these viral pathogens in dysregulating the pathways. In addition, since the molecular pathways above are multistep processes, there is need for follow up studies to interrogate the exact step or component(s) of these pathways that are targeted by the two viral pathogens. Deciphering the precise step(s) or component(s) of the MAPK and PI3K-AKT signaling pathways in cervical cells targeted by HIV-1 infection could identify potential therapeutic targets.

Our transcriptomics and proteomics analyses highlighted the regulation of the PI3K pathway. Specifically, we found increased IRS1 mRNA and protein expression from participant samples and our culture system, respectively, which is a known adaptor protein for PI3K pathway signal transduction from receptor tyrosine kinases. In addition, transcription factor activity analysis from the mRNA-seq data showed beta-Catenin to be our top hit, known to be activated by PI3K signaling. Lastly, phosphoproteomics analysis of cultured cervical cells exposed to conditioned media from HIV-1-infected T-cells confirmed activation of AKT, a protein immediately downstream of PIK3CA. Intriguingly, a recent Ugandan genomics and transcriptomics study reported that mutation and mRNA expression of PIK3CA is lower in WLWH compared to women without HIV (46). This finding suggests there to be less selective pressure for PIK3CA mutations in WLWH. In conjunction with our findings of PI3K activation from this study, we reason that HIV-1 infection may phenocopy PIK3CA pathway mutations by stimulating PI3K pathway activity through paracrine signals derived from HIV-1-infected immune cells. This concept is supported by previous studies on hepatocellular and head and neck cancers, which reported lower mutation rates of known cancer-related genes, such as FAT1, CTTNB1, TERT, TP53, in virus-infected individuals (57–59). Future studies will be required to uncover the exact factor within the immune cell secretome that activates the PI3K pathway.

In summary, our work provides valuable transcriptomic resource from an HIV-endemic area in SSA for the research community interested in studying cervical cancer in WLWH. Using systems biology approaches, we highlight the key pro-cancer pathways, such as MAPK, and PI3K-AKT, that could be underlying accelerated progression to advanced HPV-associated cervical cancer seen in WLWH. Our findings nominate these molecular pathways for follow up mechanistic studies aimed at identifying therapeutic targets for HPV-associated cervical cancer in WLWH and as therapeutic targets for WLWH.

## Materials and methods

### Recruitment and cervical sample collection

The study protocol was reviewed and approved by Moi University College of Health Sciences and Moi Teaching and Referral Hospital Institutional Ethics and Research Committee (Approval # 0003853). We also obtained a research permit from the National Commission for Science, Technology and Innovation, Kenya (Approval # NACOSTI/P/21/10523). To collect samples from health facilities within western Kenya, we obtained permission from the management of the Moi Teaching and Referral Hospital (Reference # ELD/MTRH/R&P/10/2/V.2/2010). All participants provided written informed consent prior to enrolment into the study.

In this study, we recruited participants from two Academic Model Providing Access to Healthcare (AMPATH; https://www.ampathkenya.org/) clinics in Kisumu and Trans-Nzoia counties (https://www.ampathkenya.org/where-we-work) in western Kenya. In this cross-sectional study, we targeted Black Kenyan women who were either HIV-positive or HIV-negative and were having NC, PCL or ICC based on a VIA test and/or cervical biopsy analysis. We confirmed the HIV status of the participants from their medical records. All HIV participants were on ART. Prior to the VIA test performed as detailed previously (60) by competent AMPATH nurses who are re-trained on VIA procedure every six months (61), two cervical swab samples were collected from every participant who consented and met the inclusion criteria (Table S2). Each cervical swab was put into a cryovial containing 1 mL of 1X phosphate-buffered saline (PBS) and stored at −80°C until use. Those with VIA negative and VIA positive results were classified as NC and PCL, respectively. In addition to the two cervical swabs, we collected a cervical biopsy sample from women found to be suspicious for ICC for cervical biopsy test, which we used to classify the participants into ICC group. The presence of at least one of the IARC HR-HPV types (36) in the cervical samples was assessed using Human Papillomavirus DNA Diagnostic Kit (Ref #S3057E; Sansure Biotech) following the manufacturer’s instructions.

### RNA-seq of participant cervical samples

In line with our aim of understanding the role of HIV-1 in accelerating progression to HPV-associated cervical cancer, we selected participants (n = 35) who were HR-HPV positive and HIV-infected or HIV-uninfected with NC, PCL or ICC. For RNA-seq analysis, we pooled two cervical swab samples collected per participant. Total RNA was extracted from 300 µL of the pooled cervical samples using an automated nucleic acid extraction system (TANBead maelstrom 9600) and TANBead® Nucleic Acid Extraction Kit (Ref #W665S66) following the manufacturers’ instructions. Contaminating genomic DNA was degraded using RNase-free DNase I (Thermo Scientific) following the manufacturer’s protocol. RNA was eluted in 40 µL of elution buffer. Aliquot of the RNA (2 µL) was used for bioanalysis using Agilent TapeStation system following the manufacturer’s instructions. The remaining RNA was kept at −80C until use. NEBNext® Ultra™ II RNA Library Prep Kit for Illumina® together with the NEBNext rRNA Depletion Kit version (v)2 were used for RNA library preparation as detailed in section 2 of the product manual. Libraries were processed on a SimpliAmp™ Thermal Cycler (Thermo Fisher Scientific). Sequencing libraries were quantified using Qubit™ Flex Fluorometer and Qubit Flex Assay Tube Strips after treatment with Qubit™ 1X dsDNA High Sensitivity assay kit (Invitrogen) following the manufacturer’s recommendations. Subsequently, the samples were sequenced on a Nextseq 550 platform (Illumina) using Nextseq 550 High Throughput Kit (Illumina) to generate 150 base paired end reads.

### Bioinformatics analysis of RNA-seq data

The raw RNA-seq reads were subjected to quality control using the FastQC v0.21.1 and trimmed using Cutadapt v1.18 to filter out poor-quality reads and adapters. After trimming, we mapped the reads to the human reference genome (hg38) curated by the University of California, Santa Cruz using STAR package v2.7.10b. We counted the reads per transcript using featureCount of the Subread package v2.0.6 to generate a count matrix data. We loaded the count matrix data onto RStudio v2025.05.0+496 anchored in R software v4.5.0. To identify outlier samples, we generated a PCA and inter- and intra-sample correlation plots from transformed read counts. Based on the inter and intra-sample correlation and clustering in PCA, we discarded two outliers (*i*.*e*., samples CC-21-065 and CK-21-10). We then performed differential expression analysis of the untransformed read counts using DESeq2 package v1.49.0. The DESeq2 output was visualized using various R packages (*e*.*g*., ggplot2 v3.5.2, ComplexHeatmap v2.25.0, igraph v2.1.4).

### Cell lines

For this study, we used C33A cells gifted by Jacques Archambault from McGill University, Canada. HIV-1 virus was produced using HEK293T cells (ATCC, CRL-3216). Both cells were maintained in Dulbecco’s Modified Eagle Medium (DMEM) supplemented with 10% fetal bovine serum (FBS, Gibco) and 1% penicillin/streptomycin (Corning).

### Isolation and culture of primary CD4+ T cells

Primary human CD4+ T-cells were isolated as described previously (62). In brief, PBMCs were first isolated from leukopaks (leukopaks; StemCell Technologies) from healthy deidentified donors by Ficoll centrifugation using SepMate tubes following the manufacturer’s instructions (StemCell). CD4+ T-cells were isolated from PBMCs using magnetic negative selection (StemCell, 17952) and cultured in resting-cell complete Roswell Park Memorial Institute (RPMI)-1640 medium, which was prepared by supplementing RPMI-1640 (Sigma) with 5 mM sodium pyruvate (Corning), 2 mM glutamine, 5 mM 4-(2-hydroxyethyl)-1-piperazineethanesulfonic acid (HEPES, Corning), 50 μg/mL penicillin/streptomycin (Corning), 10% FBS (Gibco), 10 IU/mL of interleukin-2 (IL-2; Miltenyi Biotec) and 5 ng/mL IL-7 (R&D Systems). The CD4+ T-cells were activated using anti-human CD3/CD8 CTS Dynabeads (Fisher Scientific #40203D) at a 1:1 cell:bead ratio at 1 × 10^6^ cells/mL.

### Preparation of HIV-1 stock

In this study, a replication-competent CXCR4-tropic subtype-B pNL4-3 (NLENG1-IRES or NLENG1I) molecular clone, in which GFP is cloned behind an internal ribosomal entry site cassette following the HIV Nef gene (63) was used. HIV stocks were prepared as described previously (62). In brief, 5 × 10^6^ HEK293T cells were transfected with 10 μg of the molecular clone (PolyJet, SignaGen) as per the manufacturer’s instructions. Day 2 and 3 post-transfection, 25 mL of the culture supernatant was harvested and filtered through 0.45 mm polyvinylidene fluoride (PVDF) filters (Millipore). The filtrate was precipitated in 8.5% polyethylene glycol (PEG, average M_n_ 6000, Sigma), 0.3 M sodium chloride at 4°C overnight. The supernatant was centrifuged at 3500 rotations per minute (rpm) for 20 min and suspended in 0.5 mL of PBS for a 100x effective concentration. Aliquots were kept at −80°C until use.

### Infection of primary CD4+ T cells with HIV-1 and isolation of secretome

Isolated CD4+ T-cells were infected with HIV-1 as described previously (62). In brief, 2 days post-isolation, CD4+ T-cells were plated in round-bottom plates in triplicate and cultured overnight in 150 μL of complete RPMI-1640. The following morning, 2.5 μL of HIV-1 stock in a 50 μL carrier volume (complete RPMI) was pipetted into each well. For the uninfected wells, 50 µL of the carrier volume without the virus was pipetted into each well. The plate was spinoculated at 1200 x g for 2 hours at 22°C to enhance infection and then washed twice with 1X PBS. Complete RPMI (*i*.*e*., 200 µL) was gently pipetted into each well and cells were incubated at 37°C in 5% CO_2_ for 3 days. The plate was then centrifuged at 400 rpm for 5 minutes at room temperature. The supernatant was transferred to a fresh plate and re-centrifuged to remove any residual cells. The supernatant was stored at −80°C awaiting stimulation of cervical cells.

### Flow cytometry analysis of primary CD4+ T cells post-HIV infection

The CD4+ T-cell pellets from the preceding section were subjected to flow cytometry analysis to assess the infection rates via GFP-tagged HIV-1. In brief, the Attune NxT Acoustic Focusing Cytometer (ThermoFisher) was used to perform flow cytometric analysis on HIV-infected and uninfected CD4+ T-cells. All events were recorded in a 100 μL sample volume following one 150 μL mixing cycle. The data in FCS3.0 file format were exported using Attune NxT Software and analyzed on FlowJo™ v10.9 using a consistent template.

### Stimulation of cervical cells with secretome of HIV-1 infected primary CD4+ T cells

Approximately 7.3 × 10^5^ C33A cells were plated into 10-cm dishes (Corning) and incubated at 37°C with 5% CO_2_ for 24 hours. The spent media was gently replaced with 12 mL of media containing 1:1 mixture of complete DMEM and supernatant collected from either HIV-infected or HIV-uninfected (controls) primary CD4+ T-cell cultures every 24 hours. The dishes were incubated at 37°C for a total of 72 hours. For each condition (*i*.*e*., HIV-infected and HIV-uninfected), three technical plates were prepared.

### Sample preparation for MS proteomics of cervical cells stimulated with secretome of HIV-1 infected primary CD4+ T-cells

Upon stimulation of cervical cells with secretome of primary CD4+ T-cells infected or uninfected with HIV-1 for 72 hours, MS analysis (*i*.*e*., abundance and phosphoproteomics) was performed on these epithelial cells as detailed previously (64). In brief, samples washed twice with cold PBS, lysed in 6LM guanidine hydrochloride (Sigma-Aldrich) and boiled at 95°C for 5 min then kept on dry ice. The lysed samples were thawed and sonicated using a probe sonicator 1x for 15 seconds at 20% amplitude. After centrifugation of the lysed samples at ~13,000 rpm for 10 minutes, supernatant was transferred to a new protein lo-bind tube. Protein was then quantified using a Bradford Assay. About 500 µg of protein sample was processed further, starting with reduction and alkylation using a 1:10 sample volume of tris-(2-carboxyethyl) (TCEP) (10 mM final) and 2-chloroacetamide (4.4mM final) for 5 minutes at 45°C with shaking. Before digestion of protein, we diluted the 6M guanidine hydrochloride 1:6 with 100 mM Tris-HCl pH8 to enhance trypsin activity and LysC proteolytic enzymes, both of which were added at a 1:100 (wt/wt) enzyme-substrate ratio and put in a water bath 37°C overnight (~16 hours). Thereafter, 10% trifluoroacetic acid (TFA) was added to each sample to attain a pH2.0. Desalting of the samples was done using a vacuum manifold with 50 mg Sep Pak C18 cartridges (Waters). Activation of each cartridge was done using 1 mL 80% acetonitrile (ACN)/0.1% TFA followed by equilibration with 3 × 1 mL of 0.1% TFA. After loading the samples, we washed the cartridges with 3 × 1 mL of 0.1% TFA. The samples were then eluted with 1 × 0.8 mL 50% ACN/0.25% formic acid (FA). Approximately 50 μg (10%) of the resulting volume was kept for abundance proteomics and the rest was used for and phosphoproteomics analysis). All samples were dried by vacuum centrifugation. For phosphopeptide enrichment of samples for phosphoproteomics, we prepared IMAC beads (Ni-NTA from Qiagen) by washing 3x with HPLC water, incubating for 30 minutes with 50 mM EDTA pH8.0, washing 3x with HPLC water, incubating with 50 mM FeCl_3_ dissolved in 10% TFA for 30 minutes at room temperature with shaking, washing 3x with and resuspending in 0.1% TFA in 80% ACN. We enriched peptides for phosphorylated peptides using a King Fisher Flex (KFF).

### Acquisition of MS proteomics data for cervical cells stimulated with secretome of HIV-1 infected primary CD4+ T-cells

Digested samples were analyzed on an Orbitrap Fusion Lumos Tribrid Mass Spectrometer (Thermo Fisher Scientific) equipped with an Easy nLC 1200 ultra-high pressure liquid chromatography system (Thermo Fisher Scientific). For both abundance and phosphoproteomics analyses, we injected samples on a C18 reverse phase column (25□cm × 75□μm packed with ReprosilPur 1.9-μm particles). We equilibrated analytical columns with 6□μL of mobile phase A with a max pressure of 650 bar. The mobile phase A consisted of 0.1% FA whereas mobile phase B consisted of 0.1% FA / 80% ACN. Peptides were separated by an organic gradient from 4% (2%) to 30% (25%) mobile phase B over 62□min followed by an increase to 45% (40%) B over 10□min, then held at 95% B for 8□min at a flow rate of 300□nL min−1. Data-independent analysis (DIA) was performed on abundance and phosphoproteomics samples using an 80-minute gradient. An MS scan at 60,000 resolving power over a scan range of 350–1100□*m*/*z*, a normalized AGC target of 300%, and an RF lens setting of 40%. This was followed by DIA scans at 15000 resolving power, using 20 *m*/*z* isolation windows over 350–1100□*m*/*z* at a normalized HCD collision energy of 30%. Loop control was set to All. To build a spectral library, one sample from each set of biological replicates was acquired in a data-dependent manner. Data-dependent analysis (DDA) was performed by acquiring a full scan over a *m*/*z* range of 350–1100 in the Orbitrap at 60,000 resolving power with a normalized AGC target of 300% and an RF lens setting of 40%. Dynamic exclusion was set to 45s, with a 10-ppm exclusion width setting. Peptides with charge states 2–6 were selected for MS/MS interrogation using higher-energy collisional dissociation (HCD), with 20 MS/MS scans per cycle. For phosphopeptide-enriched samples, we analyzed MS/MS scans in the Orbitrap using isolation width of 1.6□*m*/*z*, normalized HCD collision energy of 30%, and normalized AGC of 200% at a resolving power of 15,000 with a 22 ms and 40 ms maximum ion injection time for abundance proteomics and phosphoproteomics, respectively.

### MS proteomics data search for cervical cells

We built experiment specific libraries using mass spectra from each DDA dataset for DIA searches using the Pulsar search engine integrated into Spectronaut v19.8.250311.62635 (Huggins) by searching against a reference Uniprot Homo sapiens sequences (downloaded 7 October 2024) and an HIV-1 pNL4-3 proteome of 9 viral proteins, including gag polyprotein, pol polyprotein, vif, vpr, tat, rev, vpu, env polyprotein and nef. For protein abundance samples, data were searched using the default Biognosys (BGS) settings, variable modification of methionine oxidation, static modification of carbamidomethyl cysteine, and filtering to a final 1% FDR at the peptide, peptide spectrum match (PSM) and protein level. For phosphopeptide-enriched samples, BGS settings were modified to include phosphorylation of S, T and Y as a variable modification. We used the generated search libraries to search the DIA data. For protein abundance samples, we used default BGS settings with no data normalization. For phosphopeptide-enriched samples, we applied the PTM site localization score in Spectronaut. Imputation was not performed for all the analyses.

### MS proteomics quantitative analysis for cervical cells

Quantitative analysis and visualization were performed in the R statistical programming language (v.4.5.0). For protein abundance data, we utilized a robust R package known as Differential Enrichment analysis of Proteomics data (DEP v1.31.0), which provides an integrated analysis workflow for differential protein expression of MS proteomics data. The DEP workflow has various functions for data preparation, filtering, variance normalization (using variance stabilizing transformation; vsn) and imputation of missing values and statistical testing of differentially expressed proteins between HIV-1 infected and uninfected conditions using moderated *t*-test from limma v3.65.1. In our case, we did not impute missing values. We also minimized intra- and inter-sample variability by selecting peptides with coefficient of variation ≤0.3 for DEP workflow.

For phosphoproteomics data, we performed initial quality control analyses, such as inter-run clustering, correlations, PCA, peptide and protein counts and intensities, using R package artMS v1.27.0. Statistical analysis of phosphorylation changes between HIV-1 uninfected and infected runs were computed using peptide ion fragment data output from Spectronaut and processed using artMS. We quantified differences in phosphorylation using artMS as a wrapper around MSstats v4.17.0, via functions artMS::doSiteConversion and artMS::artmsQuantification with default settings. We grouped and quantified all peptides having the same set of phosphorylated sites into phosphorylation site groups. MSstats performs normalization by median equalization, no imputation of missing values, and median smoothing to combine intensities for multiple peptide ions or fragments into a single intensity for their phosphorylation site group. Finally, statistical tests of differences in intensity between HIV-1 infected and uninfected conditions were performed. MSstats uses the Student’s *t*-test for p-value calculation and the Benjamini–Hochberg method of FDR estimation to adjust p-values.

## Data and code availability

The mass spectrometry proteomics data have been deposited to the ProteomeXchange Consortium (http://proteomecentral.proteomexchange.org) via the PRIDE partner repository (65) with the dataset identifier PXD064529. All raw mRNA sequencing data files have been deposited in NCBI’s Gene Expression Omnibus (66) and are accessible through GEO Series accession number: GSE299125. All other data are available in the main text or the supplemental information. No custom codes were developed for this project.

## Acknowledgements

We would like to acknowledge a pilot grant from the Center for AIDS Research (CFAR; 1P30AI152501-01A1 to MB, MS, and OIF) and a Collaborative Development Award (CDA) from the HIV Accessory and Regulatory Complexes (HARC) center at the Quantitative Biosciences Institute (QBI) at UCSF (U54AI170792 to MB, MS, and OIF). This work was additionally supported by funds from a World Bank African Centres of Excellence grant (WACCBIP+NCDs: Awandare) and a DELTAS Africa II grant (DEL-22-014: Awandare), by the Wellcome Trust [DEL-15-007] and the UK Foreign, Commonwealth & Development Office, with support from the Developing Excellence in Leadership, Training and Science in Africa II (DELTAS Africa II) programme. Charles O. Olwal was supported by a WACCBIP-World Bank ACE PhD fellowship (WACCBIP+NCDs: Awandare). Mehdi Bouhaddou also would like to acknowledge the James B. Pendleton Charitable Trust and the McCarthy Family Foundation for their generous support of our research.

## Author contributions

**Conceptualization**: C.O. Olwal, G.K. Boateng, P.K. Quashie, Y. Bediako, M. Bouhaddou; **Writing-original draft**: C.O. Olwal; **Investigation**: C.O. Olwal, U. Rathore, S. Makanani, P. Kaushal, I.A. Ashley, M.R. Ummadi, V. Appiah, A.L Djomkam Zune, S. Blanc, D. Winters, Y. Delgado, M. Eckhardt, R. Kaake, D.L. Swaney; **Formal analysis**: C.O. Olwal, M. Bouhaddou; **Visualization**: C.O. Olwal; **Data curation**: C.O. Olwal, V. Appiah, A.L Djomkam Zune, D. Winters, Y. Delgado; **Supervision**: K. Muthoka, J.M. Fabius, M. Eckhardt, R. Kaake, E.O. Orang’o, D.L. Swaney, G.K. Boateng, N.J. Krogan, P.K. Quashie, Y. Bediako, M. Bouhaddou; **Funding acquisition**: N.J. Krogan, Y. Bediako, M. Bouhaddou; **Writing-review & editing**: U. Rathore, S. Makanani, P. Kaushal, I.A. Ashley, M.R. Ummadi, S. Blanc, D. Winters, Y. Delgado, K. Muthoka, J.M. Fabius, M. Eckhardt, R. Kaake, M. Su, O.I. Fregoso, J.F. Hultquist, E.O. Orang’o, D.L. Swaney, G.K. Boateng, N.J. Krogan, P.K. Quashie, Y. Bediako, M. Bouhaddou

## Competing interest

MB is a financially compensated advisor for Gen1e Life Sciences and MedStat Inc. The Krogan Laboratory has received research support from Vir Biotechnology, F Hoffmann-La Roche, and Rezo Therapeutics. NJK has a financially compensated consulting agreement with Maze Therapeutics. NJK is the President and is on the Board of Directors of Rezo Therapeutics, and he is a shareholder in Tenaya Therapeutics, Maze Therapeutics, Rezo Therapeutics, and Interline Therapeutics. The Hultquist Laboratory has received research support, paid to Northwestern University, from Gilead Sciences and is a paid consultant for Merck and Ridgeback Biotherapeutics. All the other authors have no competing interests.

## Supplementary files

**Figure S1.**
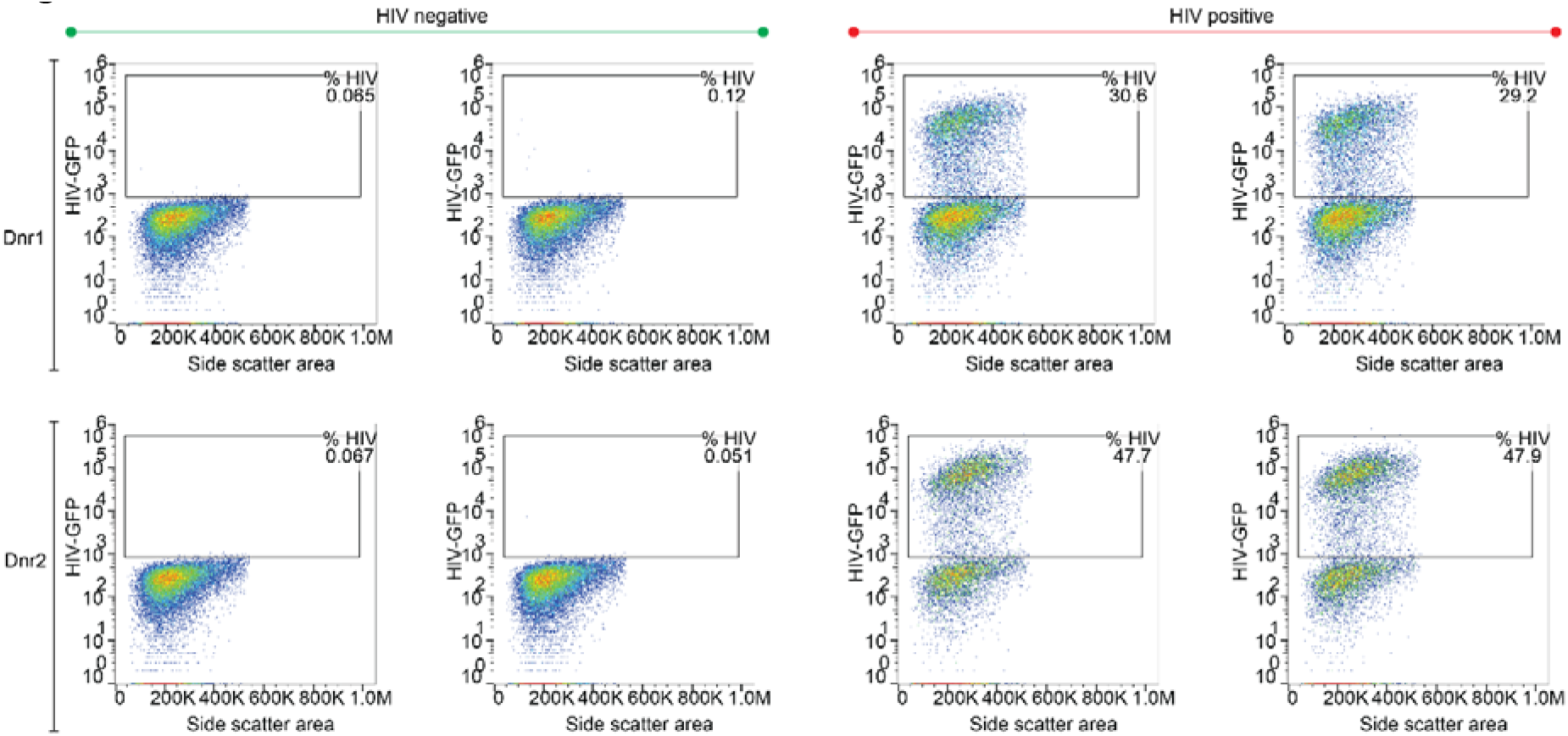
Infection of primary CD+ T-cells with HIV-1. Flow cytometry outputs for primary CD4+ T-cells uninfected (**Left**) or infected with a GFP-tagged replication competent HIV-1 strain (**Right**). The supernatants from these cultures were used to stimulate cervical cells.

**Figure S2.**
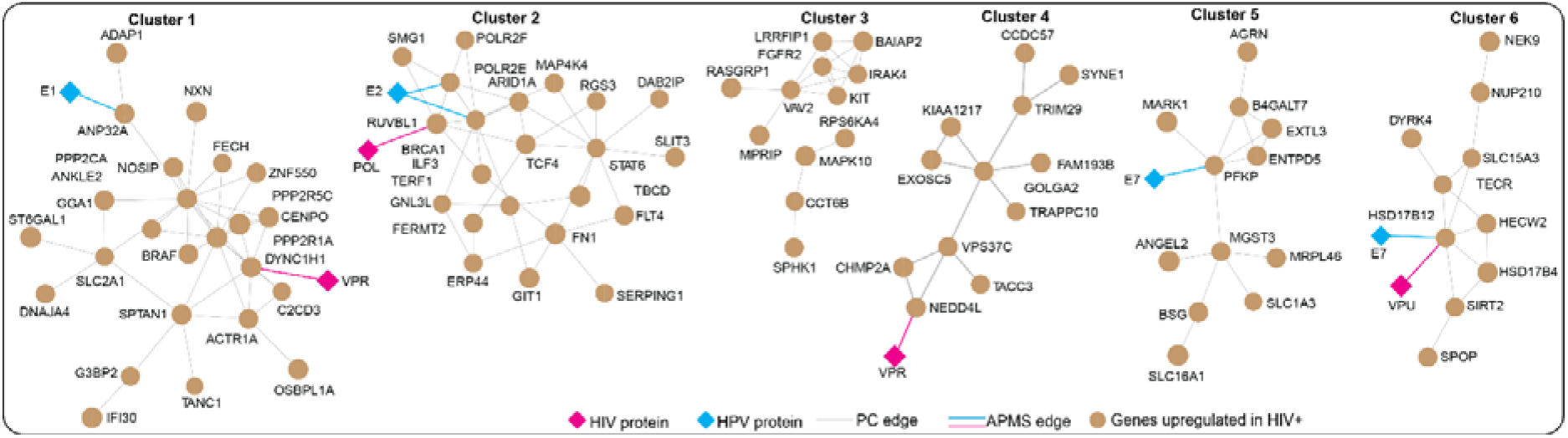
Network depicting the interconnection between genes with high mRNA over expression in our RNA-seq and TCGA cervical cancer datasets.

**Figure S3.**
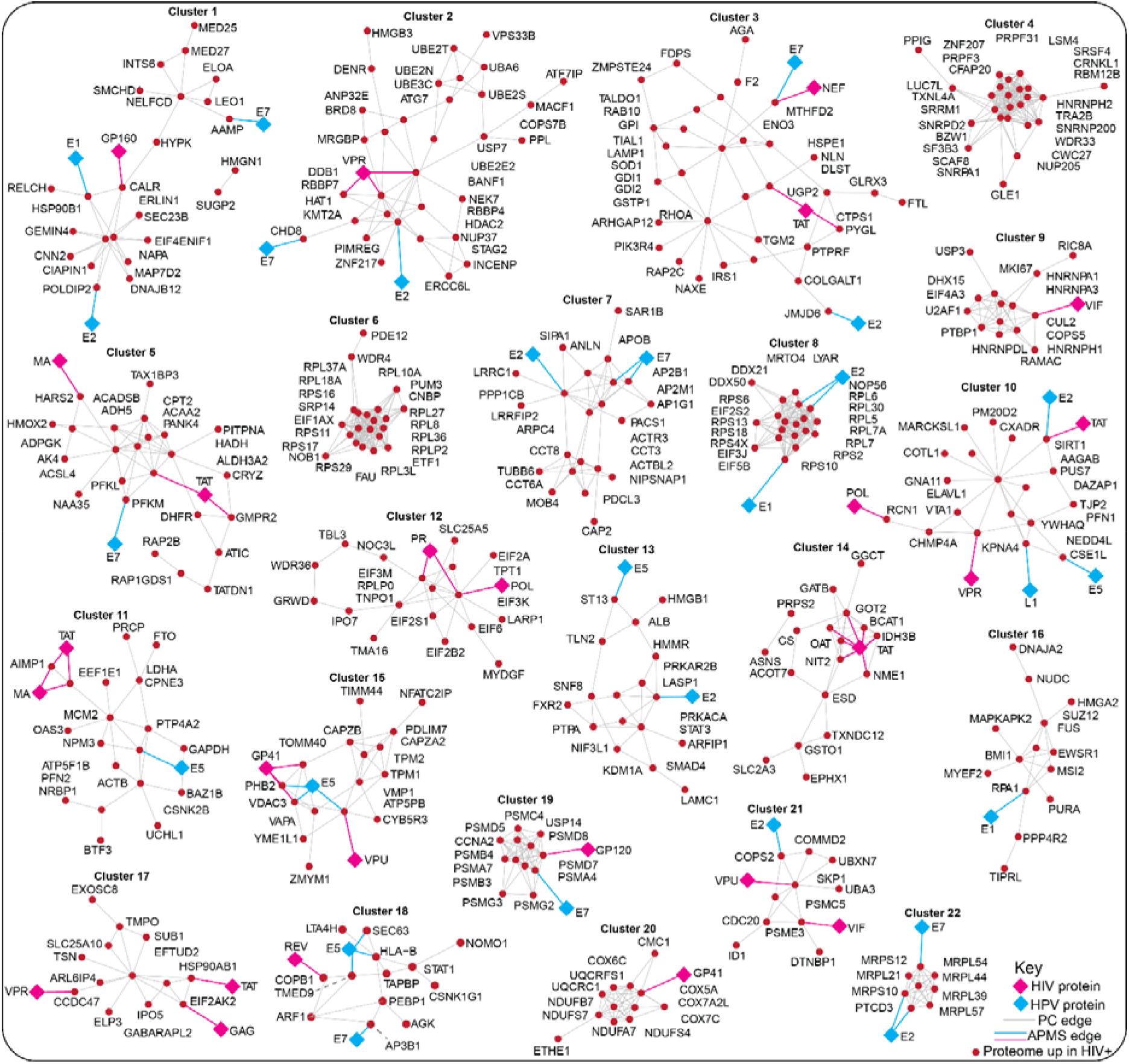
The network depicting interconnection between proteins upregulated in cervical cells exposed to secretome of primary CD4+ T-cells infected with HIV-1 relative to the uninfected controls.

**Figure S4.**
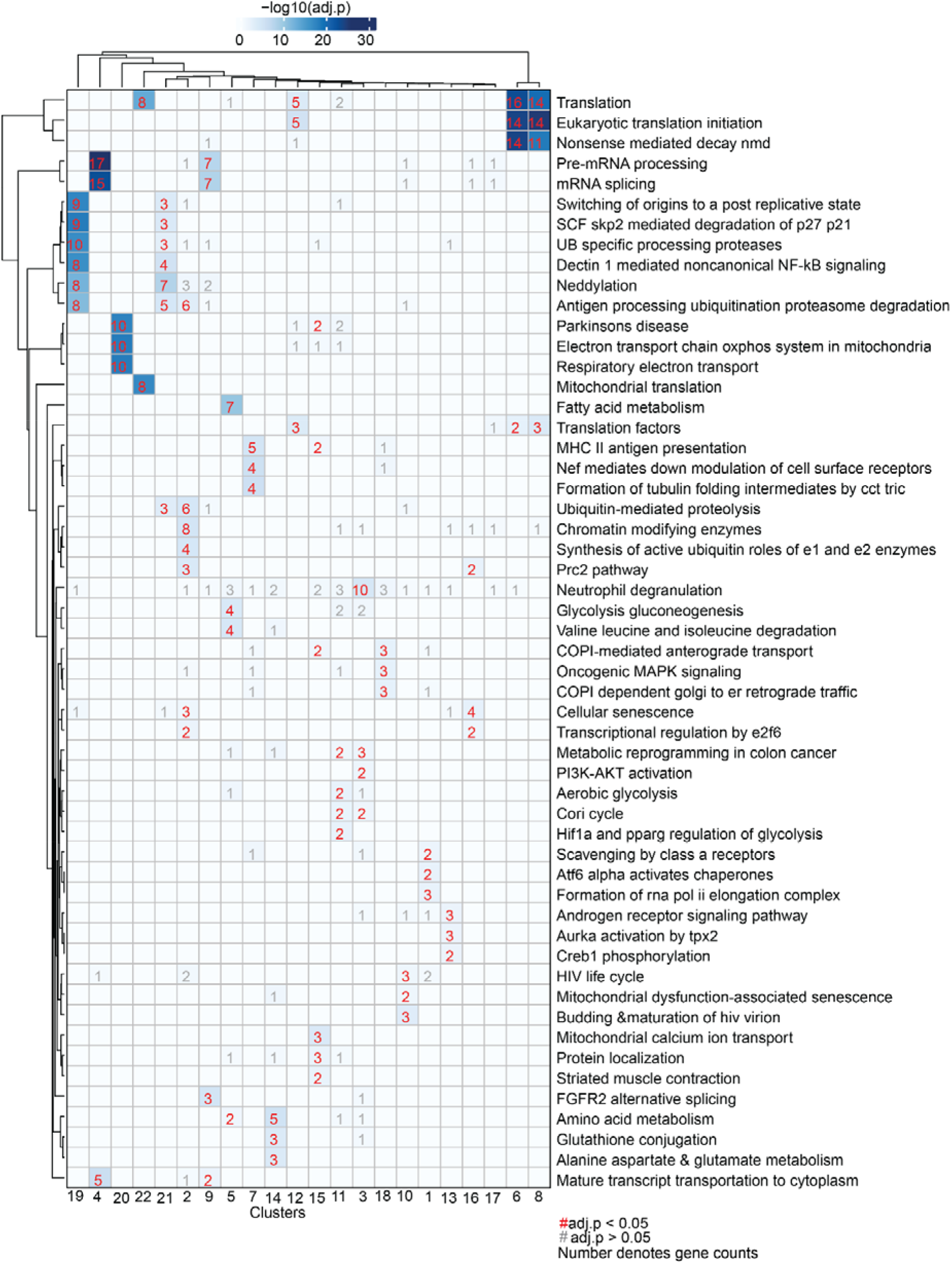
Gene set overrepresentation analysis of subnetworks generated from proteins upregulated in cervical cells treated with secretome of HIV-1 infected immune cells. Enrichment heatmap showing the canonical pathways associated with each of the subnetwork clusters in Figure S3. The gene set overrepresentation analysis was performed for the network subclusters. The p-values were calculated by hypergeometric test with multiple hypothesis testing correction (false discovery rate; FDR). A complete set of enriched biological pathways is provided in Table S4.

**Figure S5.**
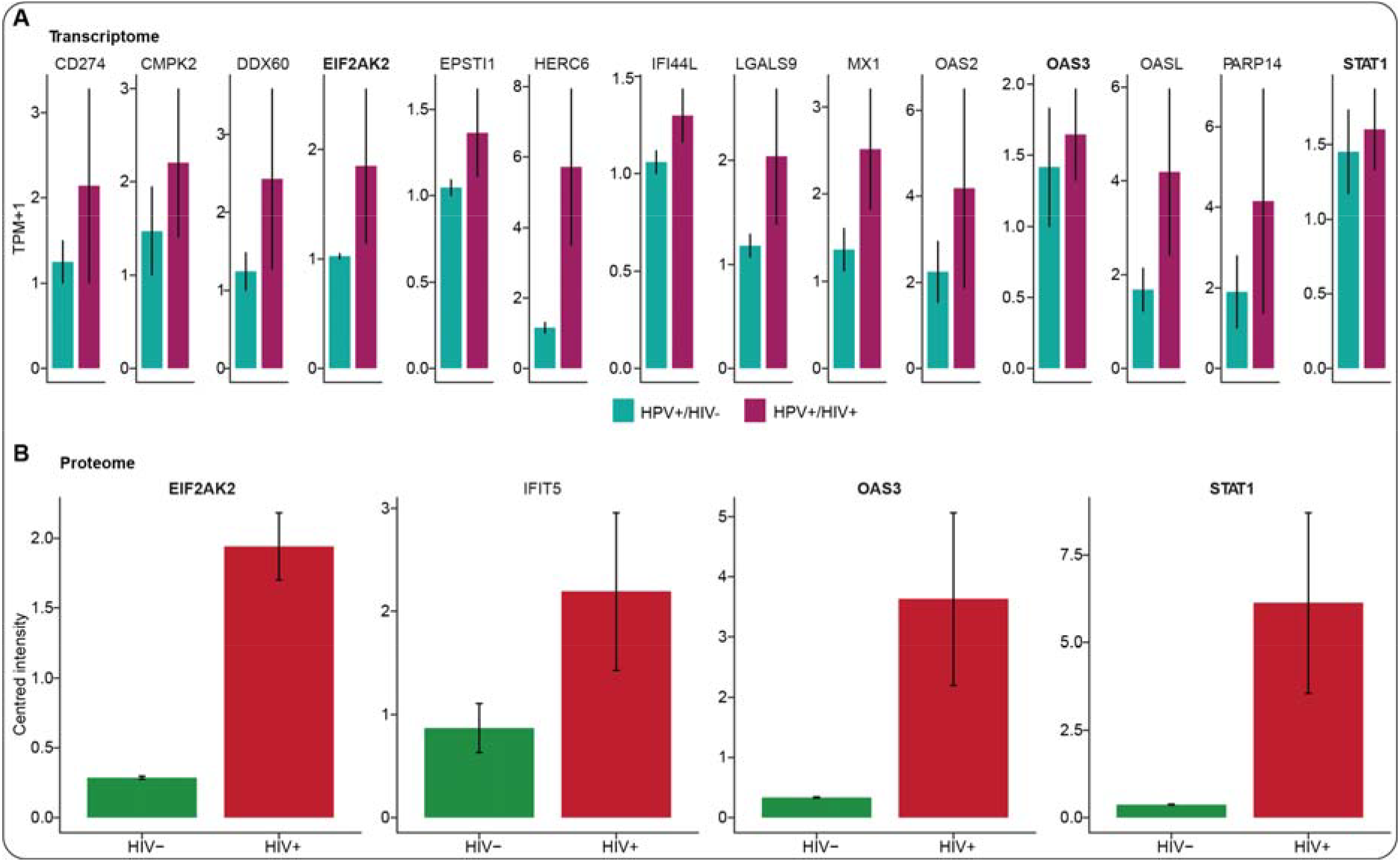
Comparison of the expression of ISGs in HIV-positive versus negative conditions. **(A)** Bar plots displaying the normalized gene expression (*i*.*e*., transcripts per million + 1) levels (mean-centered intensities) of ISGs in cervical samples of women with NC who are HIV-infected or uninfected. **(B)** Box plots showing the protein expression levels (*i*.*e*., mean-centered intensities) of known ISGs. The center line in each box depicts the median whereas the lower and upper edge of each box represent the 25th and 75th percentile values, respectively. The whiskers represent 1.5 times the interquartile range.

